# Large-scale quantitative benchmark reveals accuracy limits of proteomics

**DOI:** 10.64898/2025.12.03.692002

**Authors:** Lukasz Szyrwiel, Justus L. Grossmann, Ludwig R. Sinn, Juri Rappsilber, Vadim Demichev

## Abstract

Quantitative proteomics has become a streamlined method for high-throughput interrogation of biological samples, with a wide range of applications in both basic and clinical research. We present a benchmark data set and methodology for evaluating the accuracy of proteomics in a high-throughput setting. We build upon previous works by incorporating the variation of sample amounts and the background proteome into the experiment design. This approach allows us to comprehensively map the factors that determine the accuracy of proteomic measurements, revealing significant systematic errors missed by previous works. We envision these data to assist in future method optimisation as well as provide a comprehensive benchmark for data analysis software algorithms.

## Introduction

In recent years, mass spectrometry-based proteomics has become a routine technology for analysing biological samples. Modern liquid chromatography coupled to mass spectrometry (LC-MS) setups allow for the analysis of hundreds of samples per day, powering increasingly large experiments, further aided by robotic sample preparation, ultimately making proteomics cost-effective and broadly accessible^1^. Furthermore, advances in the instrument sensitivity have enabled a range of applications involving very low sample amounts, such as single cell proteomics^2–4^ or high-resolution spatial tissue proteomics^5,6^, as well as the analysis of low-stoichiometry post-translational modifications using limited cell numbers^7,8^.

The broad adoption of proteomics puts stringent quality control of the data in focus, in particular when it comes to applications that push the limits of LC-MS speed and sensitivity. In this work, we present a large-scale data set that enables comprehensive assessment of peptide quantification accuracy by ion mobility-resolved high-throughput mass spectrometry, covering a range of sample amounts. We build upon the LFQbench concept proposed by Navarro et al^9^, wherein tryptic digests of different organisms are mixed together in defined proportions, different across samples, with the LC-MS workflow having the goal to reconstruct the known peptide and protein ratios between samples as accurately as possible, for each species. However, our data set is different in a number of ways. First, it is acquired on a recent-generation instrument at a 200 samples per day throughput, reflecting the data quality of modern high-throughput proteomics experiments. Second, we introduce a variable proteome background (human), modelling the natural variation of the proteome of biological samples, as opposed to constant proteome of commercial tryptic digests. Third, encompassing 192 acquisitions in total, our dataset enables to evaluate how the size of the experiment affects the performance of advanced quantification algorithms incorporated in a modern proteomics data processing software, such as QuantUMS^10^ as implemented in DIA-NN^11^.

Furthermore, we envision that the new dataset can prove useful for elucidating and quantitatively characterising factors that affect quantitative precision and bias. For example, in this work, we examine how the ratios of individual fragment ions between samples are affected by the sample amount, variability in the background proteome or the predicted intensity rank of the fragment. We further establish the quantitative robustness of the fragment ion selection algorithm implemented in our DIA-NN^11^ software across different sample amounts.

## Results

### Experiment design

We aimed to establish a benchmark data set that could be used to comprehensively evaluate the accuracy and precision of quantification algorithms. To this end, we mixed tryptic digests from commercial human, yeast and *E*.*coli* whole-cell lysates in defined proportions (Fig. 1, Methods). This approach enables to evaluate how accurately the mass spectrometry method and the respective quantitative algorithms reconstruct the known analyte ratios between the mixes, for each of the species involved. In each comparison between any two mixes, we normalise the data to center *E*.*coli* analyte ratios between the mixes at 1:1 (Methods). We interpret the observed *E*.*coli* ratios following normalisation as a metric reflecting primarily the method precision, with ratios close to 1:1 being indicative of high precision. The yeast ratios, in contrast, are distinct from 1:1 and allow to evaluate the overall accuracy of the method and reveal its ratio compression, if any. Note that with 1:1 ratios any interfering signals that are consistent between the mixes act to compress the ratios, bringing them closer to 1:1 and thus improving the readout, seemingly leading to increased perceived precision. At the same time, whenever the species ratio between the mixes is distinct from 1:1, any such systematic error would be evident as ratio compression and thus would negatively affect the accuracy of the measured ratio. Together, the analysis of both *E*.*coli* and yeast ratios represents a comprehensive picture of the quantitative performance of the method. Finally, a variable human background, generated using tryptic K562 and HeLa cell line digests, serves as the source of the majority of the signals measured by mass spectrometry and thus also as the origin of most interfering signals observed in extracted ion profiles matched to *E*.*coli* and yeast peptides. Comparing a pair of mixes with the same background to a pair of mixes with distinct backgrounds allows to dissect the impact of background variation on the method performance. The background variation here is essential to better mimic the samples of interest in actual proteomics experiments, with no two samples are expected to be identical, in contrast to synthetic benchmarks.

**Figure 1.**
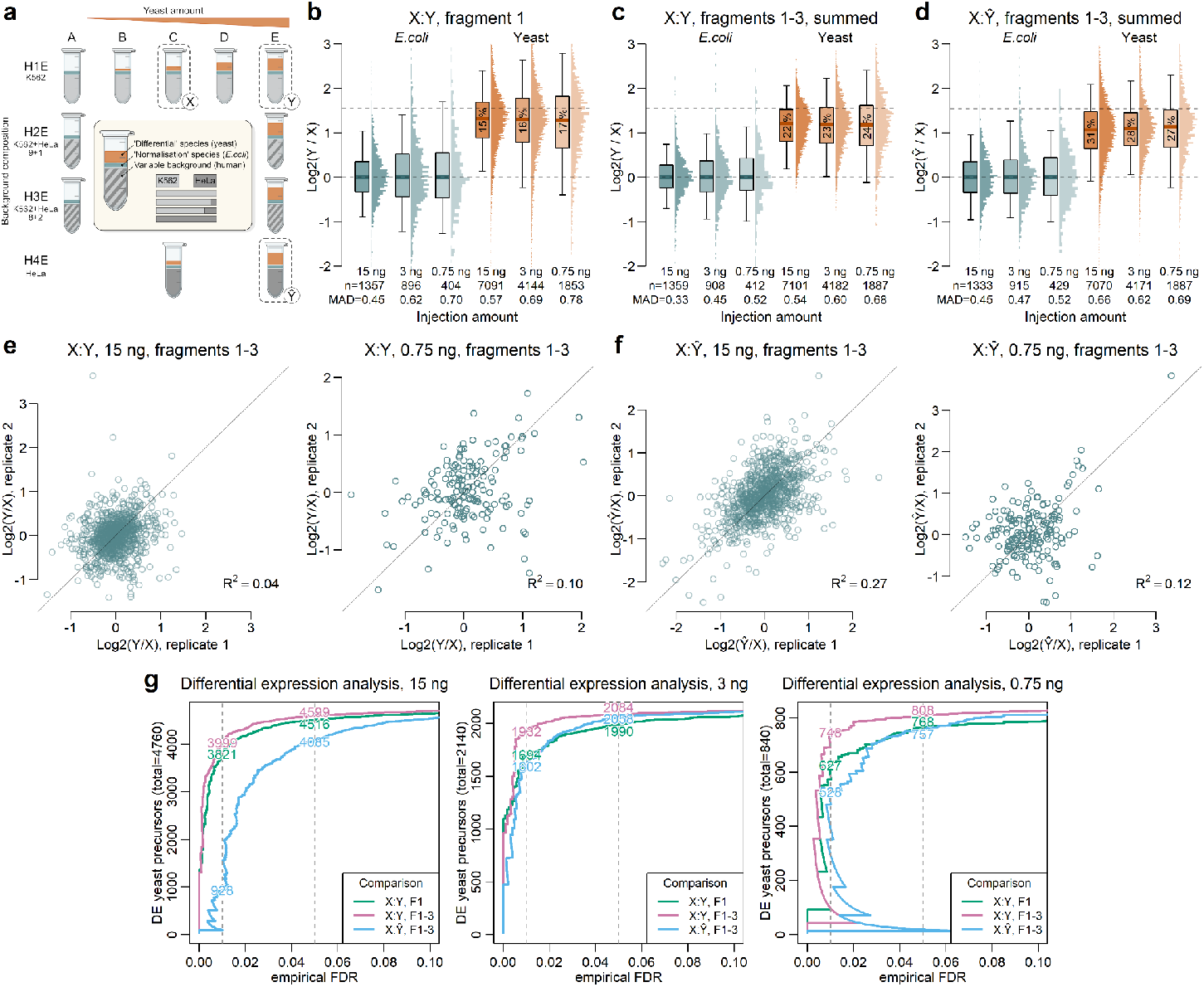
The effect of sample amount and background perturbation on quantitative precision and accuracy. **(a)** The experiment design (see Methods for the specification). Samples were prepared by mixing human tryptic proteome digests (K562 or HeLa), serving as the background, with the ‘differential’ digest (*S*.*cerevisiae* yeast) and the normalisation digest (*E*.*coli*) in defined proportions. Each sample was acquired using four loading amounts, targeting 15ng, 3ng, 1.5ng and 0.75ng of the human fraction per acquisition, each in four replicate acquisitions. An extra sample (labelled H1E-F) with an equal amount of HeLa added along with K562 as in H1E-C was further acquired (not shown on the diagram). Samples considered in this work are indicated with X, Y and Ŷ. **(b)** Quantitative ratios between X and Y across varying injection amounts, the intensity of the top library fragment used as the precursor quantity. Dotted horizontal lines indicate expected ratios. Boxes represent the interquartile range, whiskers extend to the 5%-95% quantiles. Median bias in yeast ratios is indicated in percentage points. Numbers of data points and the mean absolute deviations (MAD) from expected ratios are indicated at the bottom. **(c)** Same as (b), except the sum of the top 3 library fragments is used. **(d)** Same as (c), but comparing X and Ŷ. **(e)** Comparison of log-transformed *E*.*coli* precursor quantity ratios calculated using independent replicates, with the sum of top 3 fragments used as the precursor quantity, across 15ng and 0.75ng amounts, for the X and Y comparison. **(f)** Same for the X and Ŷ comparison. **(g)** Number of precursors vs empirical FDR (unpaired t-test, only precursors with >= 3 replicates detected in each condition) for (in different colours) comparisons in b), c) and d) - all 15ng (left plot), 3ng (middle plot) and 0.75ng (right plot).

Overall, we have acquired 192 LC-MS measurements, comprising 12 mixes, with each mix being measured at four different loads, labelled based on the approximate amount of the human sample fraction injected, respectively at 15ng, 3ng, 1.5ng and 0.75ng. Each load was acquired in four replicates. For our measurements, we used the Evosep One system coupled to timsTOF Ultra, operated in 200 SPD (Samples Per Day) mode (Methods), which has recently gained popularity for high-throughput proteomics applications. The experiment is, therefore, relevant for the wide range of arising applications of high-throughput proteomics, in particular towards analysis of low sample amounts. In the analyses below, we examine the quantitative properties of the data based on a subset of the experiment, specifically, three mixes that we label as X, Y and Ŷ, where X and Y feature the same human background (K562), distinct from the background in Ŷ (HeLa; Methods).

### Low sample amounts exasperate noise and ratio compression

First, we investigated the effect of the loaded sample amount on quantitative performance. Here, we evaluate the data from two perspectives: the overall performance across all precursor ions detected, for a given pair of mixes and the loading amount, as well as the quantitative metrics with which the same precursors are measured across different loading amounts. While the former is relevant for an overall assessment of the method, the latter reveals how specific signals are affected by a decrease in the sample load.

Naturally, we expect weaker signals to have lower signal-to-noise ratios and therefore lower precision. Indeed, we see that the variance of *E*.*coli* ratios increases at lower loads (Fig. 1b,c). However, the ratio compression, reflected by the median of yeast ratios being below the expected value, stays the same across different loads, when considering all identifications. In contrast, when considering only precursors that are detected in all conditions, the ratio compression is greater with lower loads (Supp. Fig. S1a,b). Given that integrating pure noise would result in ratios centered at 1:1, this may be explained by a greater relative contribution of random noise to low-intensity fragment quantities.

The above trends hold both when quantifying the precursor using the top library fragment, that is the fragment ion predicted in silico to be the most intense, as well as when quantifying using the sum of the top three library fragment ions. In the latter case, we observe significantly higher ratio compression at all loads, coupled to reduced variance of *E*.*coli* ratios (Fig. 1b,c). We note that such a variance reduction may be partially explained by the ratio compression itself, rather than by ‘true’ extra precision gained from integrating more fragment ion signals. However, summing multiple noisy signals, as is the case here, may also increase the overall signal-to-noise ratio and therefore improve precision.

### Fragment quantities incorporate both random and systematic errors

To investigate the sources of error in fragment ion quantities, we examined their ratios between samples obtained with either constant or variable human background. In the variable background setting, we observe a major increase in the variance of *E*.*coli* ratios at the 15ng sample amount (Fig. 1d). This effect is particularly prominent when considering the sum of the top three fragments, as opposed to just considering the top fragment (Supp. Fig. S2). Further, the variable background also resulted in increased ratio compression across all sample amounts, likely reflecting the greater overall noise levels. We next plotted the *E*.*coli* precursor ratio deviations observed in independent replicates against each other (Fig. 1e,f). Interestingly, the deviations did not correlate when considering ratios with constant background (R^2=0.04, 15ng samples), but exhibited a substantial correlation when considering a variable background (R^2=0.27, 15ng samples). This correlation was reduced but still present (R^2=0.12) when considering 0.75ng samples with a variable background. We conclude that the accuracy of fragment quantities is affected by both the random errors as well as the systematic errors caused by integrating signals that originate from interfering analytes. The use of synthetic benchmarks with variable (human) background as demonstrated here is necessary to assess the magnitude of such systematic errors.

### Full peak integration improves precision but not accuracy

In common DIA processing algorithms, peak areas for individual fragment ions are calculated by summing the observed fragment ion signals across a predefined number of scans. In the analyses above, DIA-NN reports fragment quantities obtained by integrating five consecutive scans in the vicinity of the inferred peptide elution apex. We further investigated the effect of increasing the number of summed scans to seven (Supp. Fig. S3). This led to a minimal increase in perceived precision but also resulted in a concomitant increase in ratio compression (cf. Fig. 1b). Indeed, only a minor fraction of the true fragment ion signal is detectable outside the immediate vicinity of its apex, whereas interfering signals are not expected to correlate with peptide elution, and therefore the extra signal gained by increasing the number of data points used for quantification is prone to augment the relative contribution of interferences.

### Fragment ion selection based on empirical fragment quality scores is not beneficial

Next, we investigated the effect of selecting fragment ions to be used for precursor quantification based on their quality metrics reported by DIA-NN. Specifically, DIA-NN assigns a quality score for each fragment ion of a precursor, in each run where it is identified, with this score reflecting how similar the elution profile of this fragment is to the profiles of the other fragments of the precursor. The original selection strategy proposed in DIA-NN and demonstrated to be effective when analysing SWATH data acquired with long chromatographic gradients^11^ involved selecting the top three fragments based on their average scores. We implemented this strategy also for the present dataset and observed that it had no substantial effect on either precision or accuracy of the data, compared to using the top three library fragments (Supp. Fig. S4a). We further tested if selecting fragments based on data acquired with varying loading amounts would affect performance and also did not observe any substantial differences (Supp. Fig. S4b). This indicates a high degree of robustness of this algorithm when processing highly heterogeneous data in a single experiment, such as jointly analysing single cell and bulk carrier acquisitions. Thus, while we have not observed any benefits of the original DIA-NN’s strategy here, it is also not detrimental and we speculate it may prove a more robust solution across varied LC-MS datasets, compared to the strategy based on predicted top library fragments, as the latter strongly relies on the accuracy of in silico fragment ion intensity prediction, which may suffer in the context of peptide chemical modification or changes in mass spectrometer settings.

### Differential expression numbers do not reflect quantitative accuracy

Naturally, the quantitative performance in controlled experiments with known ground truth can be evaluated by comparing the numbers of analytes identified as “differentially expressed” to the effective false discovery rate (FDR). In the context of the present experiment, every yeast precursor is differentially expressed, while every *E*.*coli* precursor is not, allowing to obtain the effective FDR at any given p-value threshold based on a t-test (Fig. 1g). Interestingly, we observed that summing the quantities of the top three fragments of a precursor resulted in slightly better differential expression detection, compared to just using the top fragment, consistent with the view that it is the precision of the measurement and not the accuracy that is the major performance determinant in this test and possibly differential expression analyses in general. Comparing the mixes with different human backgrounds (K562-only vs. K562 + HeLa), we observed significantly lower numbers of differentially regulated precursors, further supporting this view. We conclude that while such differential expression benchmarks may be useful, they cannot serve as an indicator of data quality on their own.

**Supplementary Figure S1.**
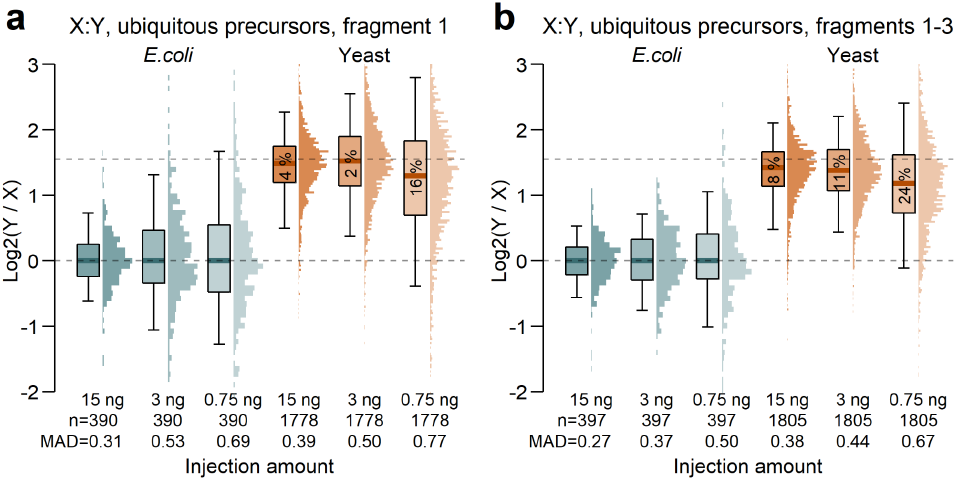
Low sample amounts are associated with loss of precision and increase in ratio compression. (a) Same as Figure 1b, except ubiquitously precursors with valid values for all conditions and amounts are shown. (b) Same as (a), except the sum of the top 3 library fragments is considered.

**Supplementary Figure S2.**
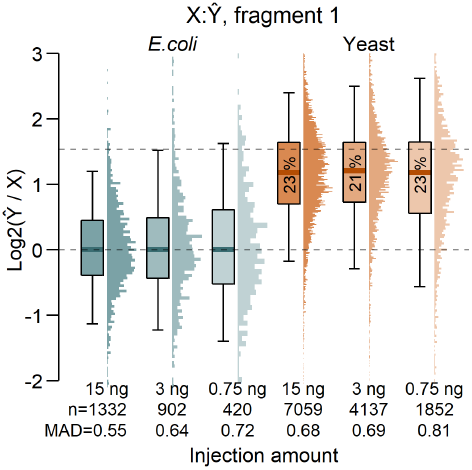
Variable background reduces measured precision. Same as Figure 1d, except only the top library fragment is considered.

**Supplementary Figure S3.**
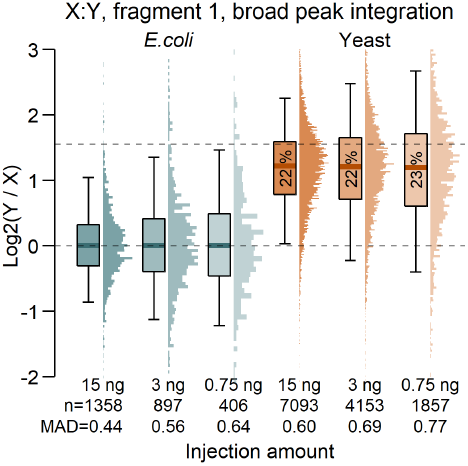
Broader peak integration reduces accuracy. Same as Figure 1b, except fragment peaks are integrated across 7 data points closest to the inferred peak apex instead of 5.

**Supplementary Figure S4.**
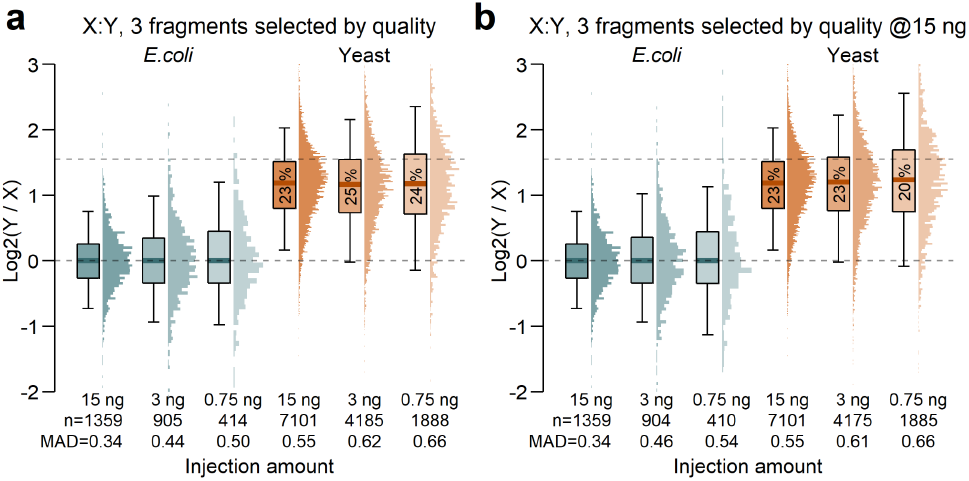
Fragment selection has no effect on precision and accuracy. (a). Same as Figure 1c, except 3 fragments are selected based on average fragment quality scores reported by DIA-NN across the respective pair of conditions. (b) Same as (a), except the fragment selection is performed solely based on 15ng acquisitions.

## Discussion

We present a benchmark and methodology for assessing the quantitative accuracy of high-throughput proteomics. The benchmark reflects how the sample amount and the resulting signal intensity modulate the quantitative precision and accuracy. Our approach also reveals the caveats of a conventional mixed-species approach with a constant background proteome^9^, wherein major systematic errors due to interfering signals of co-fragmented precursors may remain undetected, leading to optimistic precision and accuracy readouts. As expected, these interferences seem to manifest most prominently at high sample loads, where large numbers of precursor ions are present at detectable levels.

We note that our insights from the present work are limited by the instrumentation we used to acquire the data. Indeed, the time-of-flight analysers used in timsTOF-series mass spectrometers may not behave similarly to Orbitrap or Astral analysers utilised for other instruments, in particular when it comes to the ratio compression that we observe for low intensity signals here. It is more likely, however, that our observations regarding systematic errors that are detected in a variable background setting extend also to a broad range of instruments, as co-fragmentation of distinct co-eluting precursor ions is a ubiquitous characteristic of DIA mass spectrometry.

In conclusion, we expect our dataset to inform future method design on similar LC-MS setups, as well as provide a basis for evaluation of the software algorithms for precursor as well as protein quantification for DIA proteomics.

## Methods

### Sample preparation and acquisition

The tryptic digests — human (K562, Promega, V6951; HeLa, Thermo Scientific, 88329), yeast (*S. cerevisiae*, Promega, V7461), and *E. coli* (Waters, 186003196) — were mixed in the intended proportions as presented in the supplementary experiment design table provided with the manuscript. The sample preparation procedure was carefully controlled with an analytical semi-microbalance (Mettler Toledo XPR105) to ensure precise mass adjustments. The sample series were labelled according to the amount of human peptides indicated in the .d folder name (15 ng, 3 ng, 1.5 ng, and 0.75 ng). For sample handling, only Protein LoBind tubes were used, and the Evosep tips were loaded according to the standard procedure. All samples were analysed using the standard Evosep method for 200 samples per day (SPD) on the Evosep One system, equipped with a 4 cm × 150 µm column (ReproSil-Pur C18, 1.9 µm beads; Dr. Maisch, EV1107). The Evosep One was coupled to the Bruker timsTOF Ultra equipped with a Captive Spray II source (Bruker). The dia-PASEF method in high sensitivity mode was based on 100 ms accumulation and ramp time featured one MS1 and 5 MS/MS frames per cycle, each frame comprising 4 or 5 isolation windows, each 25 Th wide, resulting in a cycle time of 0.97 s. We note that after examining the data, we discovered that the sample labelled as H1E-E 1.5ng exhibits unexpected species ratios and was likely mislabelled. This sample was not included in the analyses presented here.

### Raw data processing

The raw data were processed using DIA-NN 2.2 with mass accuracies set to 15 ppm and the scan window set to 7. First, a spectral library was created using 12 acquisitions with 15ng human fraction load and then used to analyse the whole experiment with MBR disabled. To generate a separate report with fragment profiles integrated across seven data points, the --quant-wide option was passed to DIA-NN. The DIA-NN logs specifying the analysis settings have been made available (Data availability).

### Ratio analysis

For each precursor, unless indicated otherwise, the ratio between any two samples was calculated by taking the first non-zero quantity of the precursor among ordered replicate acquisitions of each sample. This way, in contrast to averaging the values across replicates, any differences in data completeness between different sample amounts would not have an effect on the perceived precision of the ratios. Given that DIA-NN would normalise its output primarily using the variable human background, which comprises the majority of precursor identifications, DIA-NN’s normalisation here does not align perfectly with the 1:1 ratio for the *E*.*coli* precursors. Therefore, all ratios were corrected via applying a shift in the log2-space to bring E.coli median ratio to 1:1 in each condition, and therefore allow the comparison of the observed yeast ratios to the known ground truth.

## Supporting information

Experiment design

## Data availability

The mass spectrometry proteomics data as well as the experiment design table have been deposited to the ProteomeXchange Consortium via the PRIDE partner repository with the dataset identifier PXD055981. The DIA-NN analysis reports and logs have been deposited to the OSF repository by the link https://osf.io/jtwvy/overview?view_only=0a100953917e41f8a826c32b3f7cd206.

## Acknowledgements

This work is funded by the German Ministry of Education and Research (BMBF), as part of the National Research Node “Mass spectrometry in Systems Medicine” (MSCoreSys), under grant agreement 161L0221.

